# Mechanistic insights into Down syndrome comorbidities via convergent RNA-seq and TWAS signals

**DOI:** 10.1101/2025.06.05.658129

**Authors:** Marc Subirana-Granés, Haoyu Zhang, Prashant Gupta, Milton Pividori

**Author notes:** Correspondence possible via GitHub Issues or email to Milton Pividori < >. msubirana · msubirana20. haoyu-zc. pragup. miltondp · miltondp · @.

## Abstract

Down syndrome (DS) is caused by trisomy of chromosome 21 and is associated with diverse clinical manifestations, yet the molecular pathways linking chromosome-21 dosage effects to DS comorbidities remain poorly defined. Here we address this gap by applying a network-based, integrative framework that combines whole-blood transcriptomic data with gene-trait associations to uncover mechanistic insights into DS-associated conditions. First, we performed matrix factorization using PLIER on Human Trisome Project (HTP) RNA-Seq profiles from 304 trisomy-21 (T21) and 95 euploid (D21) individuals, deriving 156 biologically interpretable gene modules. We then identified 92 modules whose activity differed significantly between T21 and D21 and annotated these with prior-knowledge and KEGG pathways. To connect modules to clinical traits, we integrated PrediXcan-derived TWAS results from the UK Biobank, revealing 25 T21-specific modules with significant gene-trait associations (FDR < 0.1), including modules linked to cardiovascular, hematological, immune, metabolic, and neurological phenotypes relevant to DS. Using HTP clinical records as a replication cohort, 13 of these modules reliably predicted comorbidity status (AUC > 0.65, mAPS > 0.65). Most notably module 37, an interferon-stimulated gene network, whose elevated expression robustly distinguished DS individuals with pulmonary hypertension (AUC = 0.76, mAPS = 0.73). Overall, our study demonstrates that integrating blood-derived gene modules with population-scale genetic data uncovers coherent molecular signatures underlying DS comorbidities, identifies candidate biomarkers and therapeutic targets (e.g., *ISG15, IFITs, MX1*), and highlights the power of combining transcriptomic and genetic evidence to elucidate complex disease mechanisms.

## Introduction

Down syndrome (DS) is a complex genetic disorder resulting from trisomy of human chromosome 21 (T21), characterized by a broad spectrum of clinical manifestations such as intellectual disability, congenital heart defects, pulmonary complications, metabolic abnormalities, and early-onset Alzheimer’s disease [1]. Genes on chromosome 21 typically exhibit an average 1.5-fold increase in expression [2], and thousands of genes elsewhere in the genome are also differentially expressed [3]; yet there is considerable inter-individual variability in these transcriptional perturbations [4]. Despite these widespread cis- and trans-acting dosage effects, the molecular mechanisms by which the extra chromosome orchestrates these effects remain largely undefined.

Significant advances in human genetics over the past decades have markedly improved our understanding of complex traits and diseases. Genome-wide association studies (GWAS) have identified thousands of genetic variants associated with diverse human phenotypes, underscoring the polygenic nature of most complex traits [5]. However, translating these genetic associations into biological insights has proven challenging due to the intricate interplay among multiple genes, and the fact that most GWAS variants reside within non-coding regions of the genome, complicating functional interpretations [6].

To bridge the gap between genetic associations and biological mechanisms, transcriptome-wide association studies (TWAS) integrate expression quantitative trait loci (eQTL) data to nominate candidate genes whose expression mediates trait variation [7]. In addition to revealing gene-level mechanisms, TWAS has demonstrated that many regulatory variants exhibit concordant effects across tissues, reflecting core pathways that govern transcription and splicing in diverse cell types [8,9]. Because eQTLs with broad, cross-tissue activity are more likely to impact systemic phenotypes, expression profiles from accessible tissues (for example, blood or skin) can serve as proxies for regulatory processes in less accessible organs. However, by design, TWAS typically evaluates one gene at a time, potentially overlooking critical interactions among genes within complex regulatory networks.

Recent conceptual models of complex trait genetics, particularly the omnigenic model, address these limitations by conceptualizing the genetic architecture of complex traits through highly interconnected gene regulatory networks [10]. According to this model, peripheral genes exert indirect effects by modulating core genes, which directly influence biological mechanisms. In response, machine learning-based approaches, particularly matrix factorization methods like the Pathway-Level Information ExtractoR (PLIER), have emerged as powerful tools to elucidate these regulatory networks. PLIER effectively addresses this complexity by identifying gene modules or groups of genes exhibiting coordinated expression patterns linked to defined molecular functions or pathways [11]. By decomposing high-dimensional transcriptomic data into biologically interpretable latent variables aligned with known pathways, PLIER facilitates an integrated understanding of gene regulatory mechanisms at the module level.

Building upon these methodological advancements, recent studies underscore the benefits of integrating genetic associations from GWAS and TWAS into these gene modules. Frameworks like PhenoPLIER and multimodal RNA sequencing analyses exemplified by Pantry demonstrate that combining TWAS data with module-level insights markedly enhances the interpretability of complex genetic data. This integrative strategy not only elucidates underlying disease mechanisms more effectively but also reveals novel regulatory pathways and candidate genes often overlooked by conventional analyses [12,13,14].

We framed this study within the omnigenic model, hypothesizing that the molecular mechanisms underlying DS-associated comorbidities are parallel to those observed in the general population but become altered due to regulatory cascades initiated by the extra copy of chromosome 21. This modification subsequently gives rise to the extensive clinical manifestations characteristic of DS. To test this hypothesis, we reconstructed co-expression networks from transcriptomic profiles of individuals with T21 and D21, aligning them with molecular mechanisms derived from the general population. Each resulting gene module was then systematically integrated with clinical phenotype associations obtained from large-scale population studies. By employing this integrative, network-based approach, our study aims to uncover novel insights into the precise molecular mechanisms driving phenotypic differences between individuals with T21 and euploid controls (D21).

## Results

### Overview of approach

We employed an integrative computational framework combining gene-expression and genetic-association data to elucidate molecular mechanisms underlying DS comorbidities. We first applied the PLIER matrix-decomposition method to Human Trisome Project (HTP) whole-blood RNA-Seq normalized dataset [15,16] of 304 trisomy 21 (T21) and 95 disomy 21 (D21) individuals, generating 156 biologically interpretable patterns of correlated genes termed gene modules or latent variables (LVs). Briefly, PLIER decomposed the expression matrix X into a matrix Z (gene×LV), which has gene weights for each gene module or LV, and matrix B, which quantifies the contribution of each sample to each gene module or LV. Simultaneously, the learned gene modules are aligned to a prior-knowledge matrix C of curated gene–pathway annotations, improving interpretability while reducing technical noise [11] (**Figure 1*A***)

**Figure 1:**
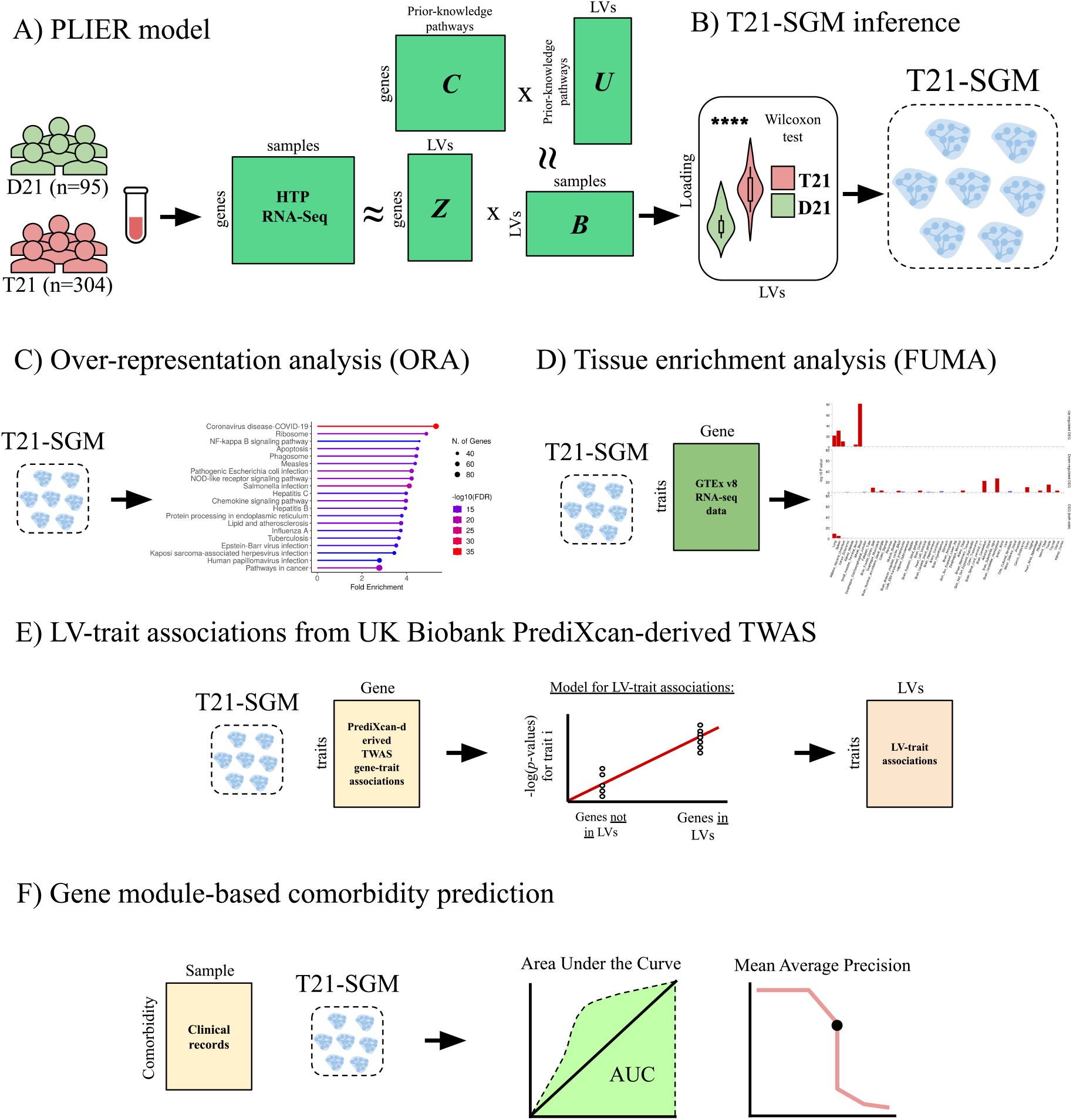
Overview of the methodology. **(A)** PLIER model. High-throughput HTP RNA-Seq data from euploid (D21, n = 95) and trisomy 21 (T21, n = 304) samples are decomposed by PLIER using prior-knowledge pathways (C) to generate gene modules/latent variables (LVs) (Z), pathway loadings (U), and sample-specific gene module loadings (B). **(B)** T21-Specific Gene Modules (T21-SGMs) inference. Wilcoxon rank-sum tests identify latent variables whose sample loadings differ significantly between T21 and D21, defining T21-SGMs. **(C)** Over-representation analysis (ORA). The top 5% of genes in each module are tested for enrichment against the KEGG pathway database. **(D)** Tissue enrichment analysis. The top 5% of genes in each module are submitted to FUMA for gene-to-tissue expression analysis using GTEx v8, identifying tissues in which module genes are significantly over-or under-expressed. **(E)** Gene module-trait association inference. For each T21-SGM, gene loadings are regressed against p-values from PrediXcan-derived TWAS in the UK Biobank (discovery cohort) to link modules with clinical phenotypes. **(F)** Module-based comorbidity prediction. Curated HTP comorbidity annotations (replication cohort) are used to evaluate each T21-SGM’s predictive performance via area under the ROC curve (AUC) and mean average precision score (mAPS).

We identified those gene modules that were differentially expressed between T21 and D21 and focused our analyses on them. For this, we then compared gene module scores derived from PLIER between T21 and D21 samples using a Wilcoxon test. Modules exhibiting statistically significant differences after multiple-testing correction were designated as T21-Specific Gene Modules (T21-SGMs) (**Figure 1*B***). To validate the molecular mechanisms inferred by PLIER, we (i) performed pathway overrepresentation analysis using the Kyoto Encyclopedia of Genes and Genomes (KEGG) database (**Figure 1*C***) and (ii) assessed tissue-specific expression patterns via the FUMA platform [17] (**Figure 1*D***).

Next, we integrate genetic data into transcriptional gene modules by connecting these gene modules with clinically relevant traits. For this, we integrated PrediXcan-derived TWAS gene-trait associations from the UK Biobank (discovery cohort). In our regression framework, each gene module is tested to determine whether genes with high loadings are strongly enriched with genes associated with a given trait [13] (**Figure 1*E***).

Finally, we leveraged clinical variables curated from the HTP dataset’s medical-record annotations (replication cohort) to assess each DS-specific module’s ability to classify comorbidity status, quantifying performance via the area under the receiver-operator characteristic curve (AUC) and the mean average precision score (mAPS). This integrated framework enabled us to systematically pinpoint gene modules with both mechanistic insight and clinical applicability in Down syndrome pathogenesis (**Figure 1*F***).

### Trisomy 21 exhibits divergent gene module profiles

To assess whether chromosome-21 triplication globally alters these transcriptomic programs, we compared module loadings between T21 and D21 groups using Wilcoxon tests, followed by Benjamini–Hochberg correction for multiple testing. Of the 156 inferred gene modules, 92 (59%) showed significantly different transcriptomic gene module loadings between karyotype groups (FDR < 0.05) (**Figure 2**). We designated these modules as T21-specific gene modules (T21-SGM). Among T21-SGM, 37 exhibited increased module activity in T21 compared to D21, whereas 55 displayed decreased activity. Furthermore, 69 out of the 92 T21-SGM (75%) were significantly aligned (FDR < 0.05, AUC > 0.6) to canonical pathways derived from prior knowledge of the general population.

**Figure 2:**
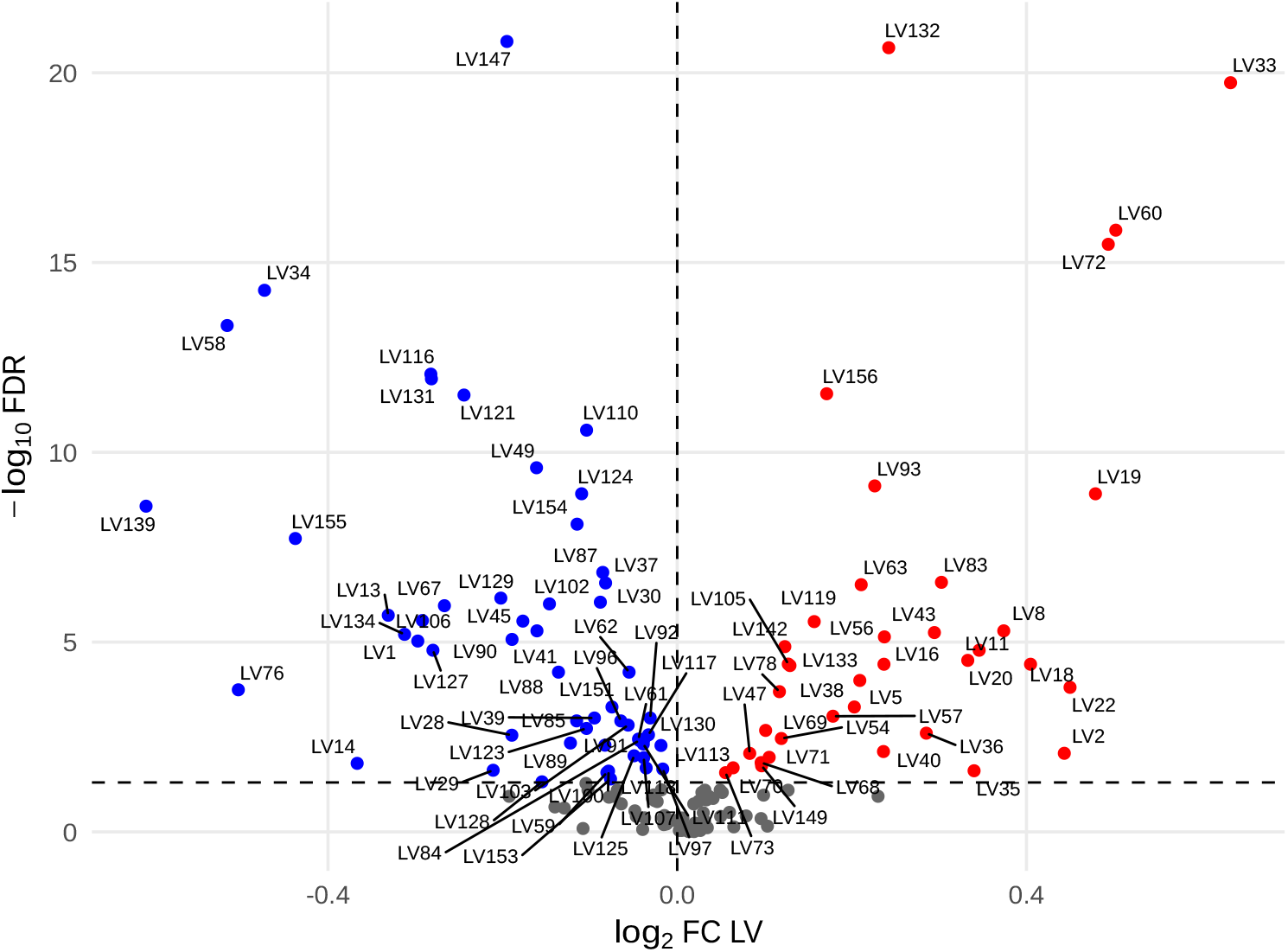
Differential activity of PLIER-derived gene modules between trisomy-21 and euploid samples. Each point represents one gene module, log_2_ fold-change in module activity (T21 vs. D21) on the x-axis and by –log10 FDR (Benjamini– Hochberg) on the y-axis. LVs significantly upregulated in T21 (FDR < 0.05, log_2_FC > 0) are shown in red; those significantly downregulated (FDR < 0.05, log_2_FC < 0) in blue; non-significant LVs (FDR ≥ 0.05) in gray. The vertical dashed line marks zero fold-change (log_2_FC = 0), and the horizontal dashed line indicates the FDR significance threshold (FDR = 0.05). A subset of the most highly perturbed T21-specific modules (e.g., LV33, LV60, LV72, LV132, LV147) is labeled.

We next examined whether the observed differences in module activity could be attributed to chromosome-21 gene dosage effects. To test this, we compared chromosome-21 gene representation between T21-SGM and non-T21-SGM. Both sets contained a comparable proportion of chromosome-21 genes (Fisher’s exact test, p-value = 0.30).

These results suggest an alteration of the transcriptional landscape in T21 rather than a shift to entirely new molecular mechanisms. These alterations occur predominantly via chromosome-21 gene overexpression rather than by overrepresentation of these genes within the affected modules.

### T21-SGMs represent tissue-specific molecular mechanisms related to DS comorbidities

Then, we sought to determine whether the molecular mechanisms driving differential gene modules in T21 align with pathways implicated in well-characterized DS comorbidities.

To this end, we identified 28 T21-SGMs exhibiting the most differential expression between T21 and euploid controls (FDR < 0.05; |log_2_FC| > 0.25). To provide a comprehensive overview of the molecular mechanisms related to these gene modules, each module was annotated through two different pathway databases: 1) PLIER’s *internal* alignment to curated gene sets during model training (FDR < 0.05; AUC > 0.6) and 2) KEGG *external* database (never used for model training) by performing pathway over-representation analysis (ORA) on the top 5 percent of module genes (FDR < 0.05).

The resulting annotations suggest a link between T21-SGMs and pathways related to DS comorbidities. Six modules were enriched for signatures of neurodegeneration, including Alzheimer’s and Parkinson’s disease pathways and broader neurodegenerative gene sets. Two converged on programmes related to acute myeloid leukemia. Nine modules were enriched for pathways associated with congenital heart defects, chronic respiratory dysfunction, adipogenesis, atherosclerosis and non-alcoholic fatty liver disease. Eleven modules indicative of immune dysregulation were enriched for interferon-mediated signaling and dendritic cell activation, while other modules reflected hematological abnormalities through signatures of myeloid and lymphoid differentiation. Some of these gene modules exhibited tissue specificity consistent with their associated pathways; for example, module 14, enriched for Alzheimer’s pathways, was overexpressed in brain tissue.

In summary, we observed that gene modules most strongly perturbed by trisomy 21 altered molecular mechanisms in the general population that can recapitulate tissue-specific transcriptional programs related to DS comorbidities.

### Integration of genetic associations from the general population provides mechanistic insights into DS comorbidities

Motivated by the observation that T21-SGMs recapitulate molecular pathways underlying well-characterized DS comorbidities, we tested whether these transcriptomic signatures also capture phenotypic associations of DS-relevant conditions. However, a major limitation of our approach arises from using peripheral whole-blood samples to infer molecular mechanisms operative in tissues directly affected by DS comorbidities. Although whole blood is readily accessible, it may not fully represent tissue-specific gene regulation dynamics. Previous studies using TWAS integrating PrediXcan and GTEx data have demonstrated that many regulatory variants exhibit conserved effects across tissues, thereby enabling the inference of biological mechanisms in inaccessible tissues using gene expression profiles from accessible tissues like blood [8,9]. Thus, integrating TWAS-derived gene-trait associations partially mitigates this limitation by enabling the identification of tissue-specific regulatory effects relevant to DS comorbidities.

To this end, we integrated PrediXcan-derived transcriptome-wide association study (TWAS) results from the UK Biobank (using diseases with ≥1,000 cases and quantitative traits with ≥5,000 samples) with our 28 T21-SGMs (discovery cohort). For each trait, we regressed its TWAS p-values against a module’s gene loadings, testing whether genes that strongly belong to the gene module are also more strongly associated with a given trait than the rest of the genes [13].

This analysis identified 25 gene modules exhibiting significant trait associations, yielding 109 module–trait links at an FDR < 0.1, of which 63 correspond to clinical phenotypes with established relevance to DS [1]. These associations map onto eleven core comorbidity categories: two modules associated with psychiatric traits (general happiness; methods of self-harm; self-harm in the past year), three modules associated with cardiovascular traits (atherosclerotic heart disease; doxazosin, an α-blocker used to treat hypertension; Adipine MR 10 m/r tablet, a calcium-channel blocker for hypertension), eleven modules associated with hematological disorders (lymphocyte count; eosinophil count; monocyte count; neutrophil count; basophil count; erythrocyte count; leukocyte count; reticulocyte count; platelet count), two modules associated with immune dysfunction (sulfasalazine used in autoimmune and inflammatory conditions; Mucodyne, a mucolytic agent often used during infection or inflammation), one module associated with obesity (weight), three modules associated with bowel dysfunction and gastrointestinal structural defects (ulcerative colitis; peritonitis; Colofac-100 tablet, prescribed for irritable bowel syndrome), one module associated with refractive errors (astigmatism), three modules associated with respiratory diseases (asthma; Symbicort 100/6 Turbohaler, a combination inhaler used in asthma management; prednisolone, a systemic corticosteroid frequently used in asthma and other respiratory conditions), two modules associated with type 1 diabetes (type 1 diabetes mellitus; diabetic ketoacidosis; insulin), six modules associated with autoimmune skin conditions (psoriasis; Dovobet ointment, a topical treatment for psoriasis; Dovonex cream, a vitamin D analog for psoriasis; eczema/dermatitis; hay fever; allergic rhinitis or eczema), and five modules associated with other autoimmune disorders (rheumatoid arthritis; polyarthropathies; methotrexate used in autoimmune diseases; folic acid co-administered with methotrexate; sulfasalazine used in autoimmune gastrointestinal and joint diseases).

Collectively, these results demonstrate that T21-specific gene modules not only capture transcriptomic molecular mechanisms related to T21 via pathway analyses but also recapitulate DS clinical phenotypes via genetic studies and trait associations, indicating that this methodology can infer mechanistic insights into DS co-occurring conditions.

### Gene modules predict comorbidity status in Down syndrome

To evaluate whether T21-SGMs recapitulate discovery cohort clinical comorbidities and thereby infer mechanistic insights into DS co-occurring conditions, we used the HTP dataset’s curated medical-record annotations as a replication cohort and quantified each module’s ability to predict comorbidity status. For every T21-SGM, we used matrix B (samples x LVs) to assess whether a gene module was overrepresented with samples from a particular DS comorbidity. For this, we computed the area under the receiver operating characteristic curve (AUC) and the mean average precision score (mAPS). We identified 13 T21-SGMs capable of predicting comorbidity status with high performance (AUC > 0.65, mAPS > 0.65) (**Figure 3**). Several of these gene modules demonstrate significant trait associations identified in the discovery UK Biobank cohort through integrated genetic analyses, uncover underlying molecular mechanisms via pathway enrichment, and display tissue-specific enrichment concordant with predicted comorbidities based on discovery HTP metadata. This convergence of different sources of evidence prioritizes potential mechanistic links between the molecular features captured by the gene modules and the manifestation of co-occurring conditions in individuals with DS.

**Figure 3:**
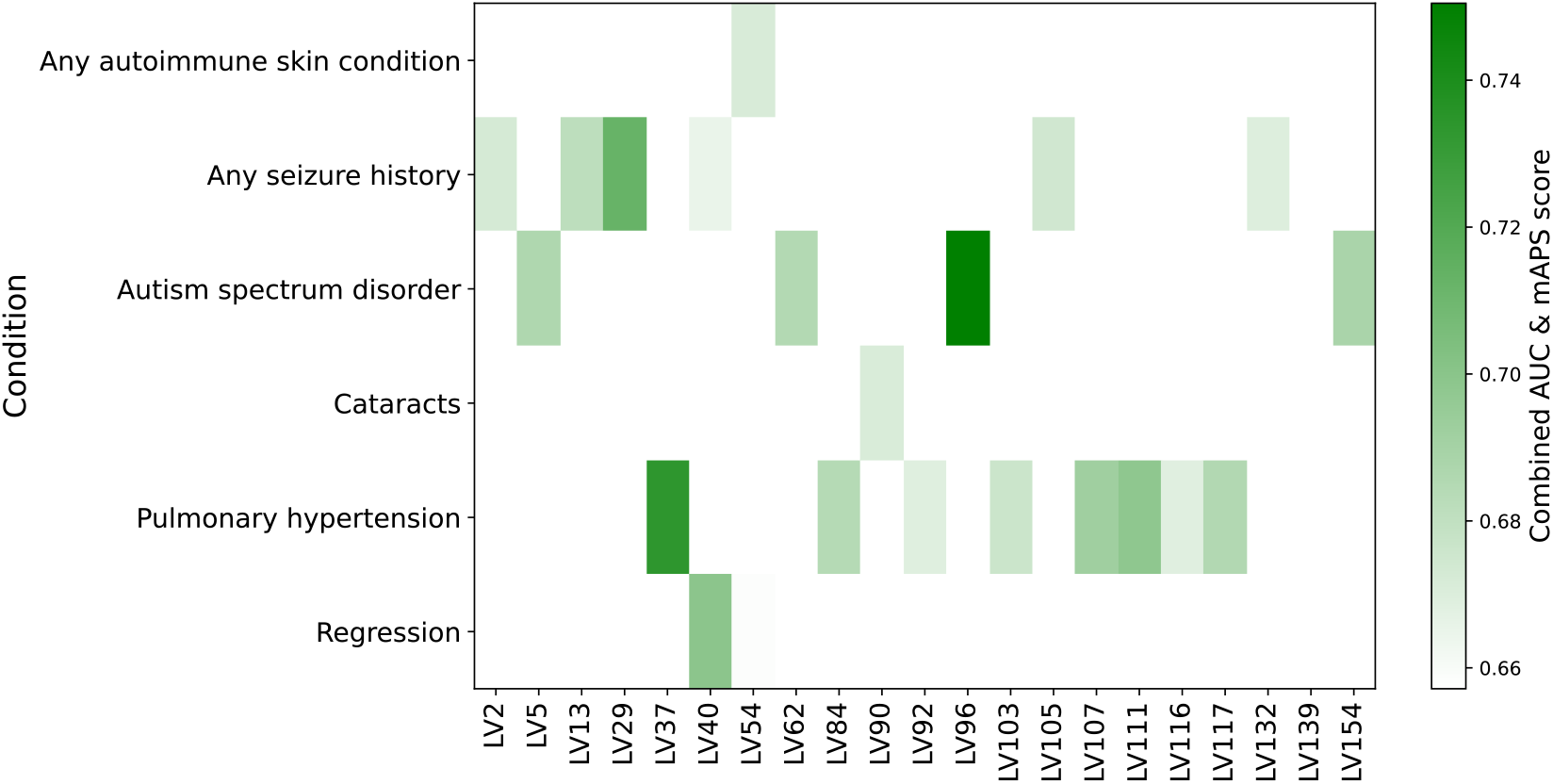
Predictive performance of T21-specific gene modules for Down syndrome comorbidities. The heatmap describes the combined predictive score (the average of AUC and mAPS) for each gene module that exceeded both AUC > 0.65 and mAPS > 0.65 when discriminating comorbidity status among T21 individuals. Columns represent individual gene modules and rows correspond to DS comorbidities from HTP replication clinical records. The color scale reflects the combined-score values, indicating joint predictive accuracy.

### Gene module 37 captures an interferon-stimulated gene signature driving pulmonary hypertension in Down Syndrome

We systematically examined sources of evidence (enriched pathways, tissue specificity, module-trait associations, and clinical metadata) to establish associations between DS comorbidities and gene modules.

All sources of evidence consistently indicate that gene module 37 is robustly associated with pulmonary hypertension (PH) in individuals with DS. Gene module 37 expression levels are significantly higher in T21 subjects with PH compared to those without PH (Student’s t-test, p-value < 0.01) (**Figure 4*A***). AUC and mAPS methodologies indicated that gene module 37 significantly differentiated between samples with and without pulmonary hypertension (AUC=0.76 and mAPS=0.73) (**Figure 4*B***), and samples within the top quartile (≥75th percentile) of expression for these gene modules showed a significant overrepresentation of pulmonary hypertension cases (p-value < 0.01).

**Figure 4:**
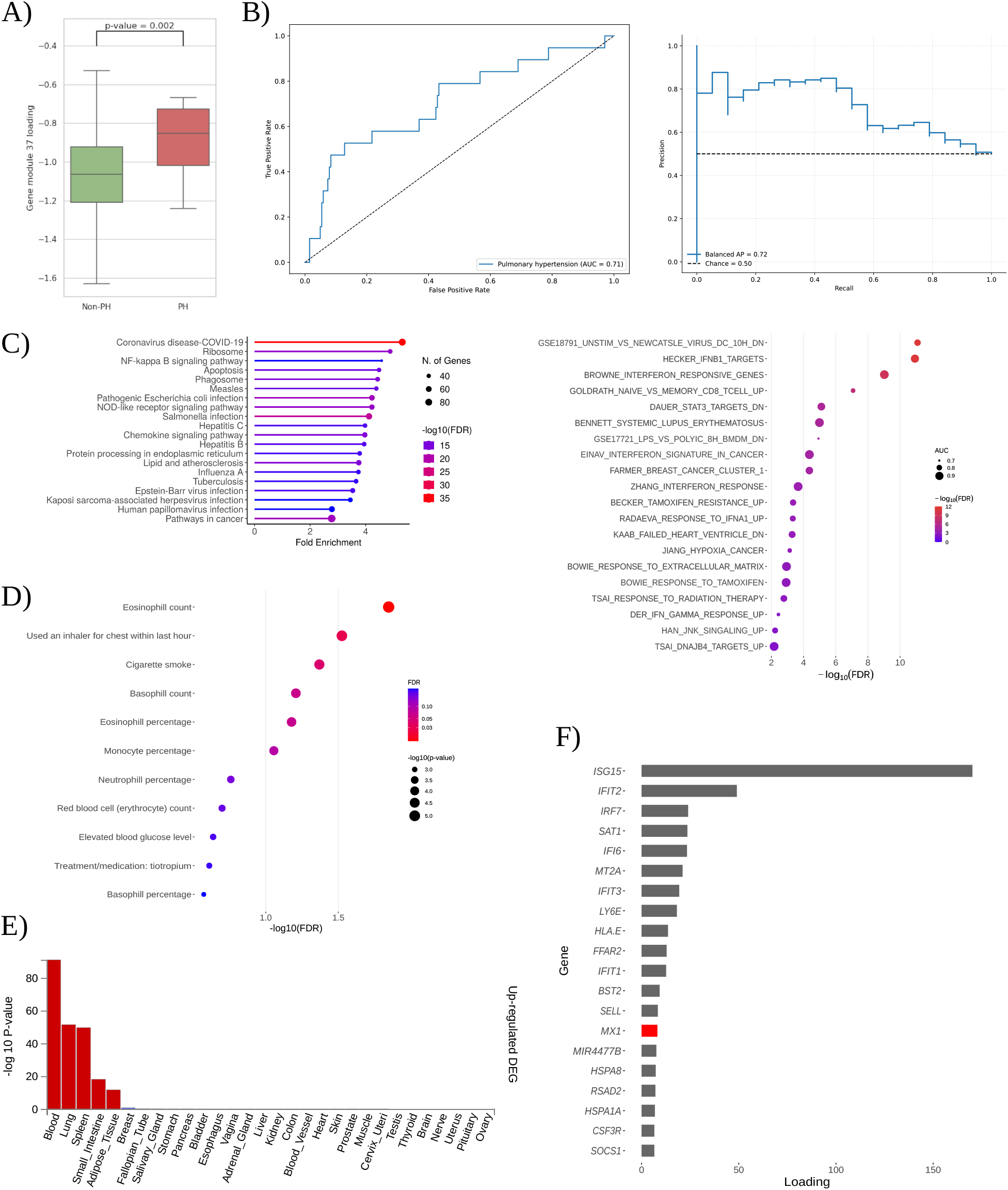
Comprehensive characterization of gene module 37 in Down syndrome–associated pulmonary hypertension. **(A)** Gene module 37 expression in T21 individuals without pulmonary hypertension (Non-PH) versus with pulmonary hypertension (PH); Student’s t-test p-value = 0.002 is indicated. **(B)** Classification performance of gene module 37 for PH: receiver-operating-characteristic curve (left) and precision-recall curve (right). **(C)** Molecular mechanisms inference for gene model 37. Upper: KEGG over-representation analysis of gene module 37’s top 5 % genes, showing fold enrichment (x-axis), –log10 FDR (color), and gene set size (point size). Lower: PLIER prior-knowledge alignment of gene module 37, plotting curated gene sets by AUC (point size) and –log10 FDR (color). **(D)** Gene module-trait association from PrediXcan-derived TWAS associations for gene module 37. Each bubble represents a clinical trait, sized by number of cases and colored by – log10 FDR. **(E)** FUMA tissue specificity results for gene module 37 genes, showing –log10 p-values across GTEx tissues. **(F)** Gene-loading profile for gene module 37 of the top 20 genes. Those encoded on chromosome 21 are highlighted in red.

Pathway enrichment analyses strongly support gene module 37’s relevance to PH. PLIER identified significant enrichment (FDR < 0.05; AUC > 0.6) in pathways such as STAT3 and JNK signaling, both implicated in pulmonary arterial hypertension (PAH) pathogenesis [18,19]. Similarly, gene expression patterns in heart failure associated with PH were enriched [20]. Independently, overrepresentation analysis (ORA) of gene module 37’s top genes revealed significant enrichment (FDR < 0.05) in NF-κB and NOD-like receptor signaling pathways, both directly linked to PAH [21,22,23]. Additional enrichment for COVID-19 and lipid/atherosclerosis pathways highlights potential inflammatory and metabolic contributions to PH [24] (**Figure 4*C***).

Tissue-specific expression analysis confirmed that lung, spleen, and whole blood are the primary sites of module activity (FDR < 0.05; |log_2_ fold change| ≥ 0.58), aligning with known tissues affected by PH (**Figure 4*D***).

Module-trait association analyses from the UK Biobank (discovery cohort) further corroborate gene module 37’s role in PH by revealing significant links (FDR < 0.1) to clinical traits, including tiotropium treatment for chronic obstructive pulmonary disease (COPD), elevated blood glucose, recent inhaler use, and cigarette smoke exposure [25,26,27,28] (**Figure 4*E***). These traits highlight clinically relevant factors and environmental risk contributors to PH.

At the individual gene level, gene module 37 prominently features interferon-stimulated genes (ISGs), including *ISG15, IFIT1-3, RSAD2*, and *IRF7* (**Figure 4*F***). Many of these genes have established roles in PH pathology. *ISG15* is identified as both a biomarker and mediator of pulmonary vascular remodeling [29,30]. *IFIT* family genes form part of a characteristic transcriptomic signature in PAH patients [31,32], while *IRF7* is upregulated in hypoxia-induced rodent models of PH [33]. *RSAD2* expression is elevated in peripheral neutrophils of idiopathic PAH patients, indicative of enhanced interferon-mediated inflammation [30].

Gene module 37 notably includes *MX1*, ranking as the 14th top gene. *MX1* is located on chromosome 21 and encodes a dynamin-like GTPase with antiviral functions, although its role in DS pathogenesis remains unclear. Recent studies implicate *MX1* in AP-1 transcriptional regulation in DS [34]. AP-1 itself plays a crucial role in pulmonary hypertension, driving aberrant pulmonary artery smooth muscle cell proliferation via endothelin-1 signaling in idiopathic pulmonary arterial hypertension (IPAH) patients [35]. These findings suggest that *MX1* may interact with AP–1–mediated pathways in pulmonary vasculature, highlighting *MX1* as a potential therapeutic target for pulmonary hypertension in DS.

Collectively, these multiple lines of evidence converge to support gene module 37’s critical role in PH within DS, identifying specific genes and pathways as promising biomarkers and therapeutic targets.

## Discussion

Despite substantial progress in understanding the genetic basis of DS, elucidating how trisomy of chromosome 21 broadly disrupts molecular networks, leading to its diverse co-ocurrent conditions, remains challenging. Our study addresses this crucial gap by integrating transcriptomic and genetic data to unravel the molecular underpinnings of DS comorbidities. Based on the omnigenic model, we proposed that chromosome 21 trisomy initiates regulatory cascades affecting gene networks analogous to those observed in the general population. Leveraging PLIER-derived gene modules, TWAS gene-trait associations, and clinical data from the HTP records, we systematically connected transcriptomic disruptions with DS-associated phenotypes. The identification of trisomy 21-specific gene modules, significantly differentially expressed compared to euploid controls, highlights potential causal pathways driving DS comorbidities. Notably, these modules predominantly align with canonical pathways rather than entirely novel mechanisms, implicated in DS comorbidities, including neurodegeneration, cardiovascular dysfunction, hematological abnormalities, and immune perturbations.

A significant strength of our analysis lies in overcoming the limitation of using whole blood RNA-seq data by integrating diverse datasets. Previous research supports that many cis-regulatory genetic variants have conserved effects across multiple tissues, reinforcing the existence of shared regulatory pathways [8,9,36]. Consistent with these findings, our integrative methodology demonstrates that transcriptomic blood-derived gene modules significantly reflect mechanisms in tissues directly implicated in DS comorbidities. T21-SGM significantly overlap GTEx signatures in lung, brain, and immune tissues implicated in DS comorbidities. TWAS-derived gene-trait associations link these modules to eleven core DS comorbidity categories (e.g., psychiatric traits, cardiovascular conditions, hematological measures, immune dysfunction, obesity, respiratory diseases, autoimmune disorders), capturing broad regulatory dynamics across different biological contexts.

Ultimately, our integrative framework bridges transcriptomic signatures with genetic associations, providing deeper insights into DS-related complex diseases. While recognizing that future studies using single-cell RNA-seq from affected tissues and rigorous experimental validations in cellular or animal models are essential, our findings substantially advance the understanding of DS comorbidities. The identification of clinically relevant gene modules, exemplified by module 37’s robust association with PH, underscores their potential utility as biomarkers and therapeutic targets. Specifically, the enrichment of interferon-stimulated genes in this module identifies and prioritize promising therapeutic candidates, such as *ISG15, IFIT* family genes, and chromosome-21 gene *MX1*. Framed within the omnigenic model, our results suggest that chromosomal dosage imbalance initiates regulatory cascades, highlighting novel intervention points for treating DS comorbidities.

## Methods

### Cohort and data preprocessing

This study was conducted under a protocol approved by the Colorado Multiple Institutional Review Board (COMIRB #15–2170). Participants were enrolled through the Crnic Institute’s Human Trisome Project (HTP; www.trisome.org). Demographic and clinical metadata were collected via structured questionnaires administered to participants and caregivers and supplemented by abstraction of the medical record.

Peripheral whole-blood samples were drawn into PAXgene RNA Tubes (Qiagen) from 304 individuals with trisomy 21 (T21) and 95 euploid controls (D21). Total RNA was extracted and assessed for integrity according to established HTP protocols [2,15]. Sequencing libraries were generated and sequenced on an Illumina NovaSeq 6000 instrument (Novogene) with paired-end reads. Raw sequence data were quality-filtered and adapter-trimmed, then aligned to the GRCh38 human reference genome using STAR [37]. Gene-level expression values were computed as transcripts per million (TPM).

### PLIER HTP model generation

For the HTP model generation, we applied the PLIER algorithm (PLIER R package v0.1.6; https://github.com/wgmao/PLIER) to the z-score-normalized TPM HTP RNA-Seq expression matrix (36,362 genes x 399 samples) [11]. Briefly, the gene-by-sample TPM expression matrix (g × s) was first z-score normalized on a per-gene basis using the rowNorm function. PLIER then factorizes this matrix into a gene loadings matrix Z (g × k) and a latent activity matrix B (k × s), where k denotes the number of latent variables. To guide this decomposition, we supplied prior knowledge in the form of a binary gene-by-geneset membership matrix C (g × p), comprising MSigDB v4.0 C2 (curated gene sets), C6 (oncogenic signatures), C7 (immunologic signatures), bloodCellMarkersIRISDMAP, and svmMarkers. During optimization, PLIER enforces correspondence between Z and its pathway-based prediction (C·U, with U a p × k coefficient matrix) by penalizing the distance between these matrices, and applies an elastic-net penalty on U to ensure that each latent factor represents a sparse subset of pathways. The joint objective is solved end-to-end via block coordinate minimization. We determined k by applying the num.pcs function to the normalized TPM matrix, yielding 156 significant components. Model fitting used the k=156 and frac=0.7 options. L1 and L2 regularization parameters were inferred automatically. Upon convergence, both the Z and B matrices were mean-centered and scaled to unit variance to facilitate downstream comparisons and integrative analyses.

### Regression model for latent variable-trait associations

We inferred gene-trait associations from the PrediXcan methods with PLIER-derived latent variables using generalized least squares (GLS) regression, described in detail in [13]. Briefly, PrediXcan methods included S-PrediXcan (gene–tissue associations) and S-MultiXcan (cross-tissue meta-analysis). For each latent variable l, we constructed a binary indicator vector sı that flags genes within the top 1% of absolute loadings within that LV. In the GLS framework, the vector of S-MultiXcan p-values for a given trait was regressed on this indicator, together with an intercept and gene-specific covariates, while modeling residuals with a covariance structure given by the gene–gene correlation matrix. This approach tests whether genes most heavily weighted in a latent variable show stronger trait associations than other genes.

### Pathway over-Representation analysis (ORA)

For each latent variable, we extracted the top 5 % of genes ranked by absolute loading values and performed over-representation analysis against the Kyoto Encyclopedia of Genes and Genomes (KEGG) database using the clusterProfiler R package (v4.0) [38]. The universe for enrichment was defined as all genes quantified in the HTP RNA-seq dataset. Resulting p-values were adjusted for multiple testing via the Benjamini–Hochberg procedure, and pathways with an adjusted p-value < 0.05 were considered significantly enriched.

### Tissue specificity annotation

Tissue-specific expression of LV-associated genes was annotated using the FUMA web platform (v1.8.0; accessed May 26, 2025) [17]. The goal of this analysis was to test whether LVs found in our whole blood RNA-seq dataset from HTP also resembled mechanisms preserved in other tissues. For each latent variable, its constituent genes were submitted to FUMA’s Tissue Expression Analysis module against GTEx v8. Default parameters were applied to test for significant over-expression of each gene set across 54 GTEx tissues; tissues meeting FUMA’s built-in significance threshold were reported as showing LV-specific expression (Bonferroni-corrected P < 0.05 and |log_2_ fold change| ≥ 0.58).

### Comorbidity classification performance

We assessed each trisomy-21-specific LV’s ability to predict between T21 individuals with and without a given comorbidity using two complementary metrics. First, for each annotated comorbidity, we retrieved the subset of T21 participants classified as cases or controls and extracted their corresponding LV activity scores. Discriminative performance was quantified by the area under the receiver operating characteristic curve (ROC AUC), computed with scikit-learn’s (v1.6.1) [39] roc_auc_score function, and by the mean average precision score (mAPS). Second, the mAPS was calculated using the average_precision_score function and further balanced by repeatedly subsampling the negative class to match the number of cases (50 iterations) and averaging the resulting precision scores. Only LVs meeting all three criteria, ROC AUC > 0.65 and mAPS > 0.65 were included.

### Data and code availability

All original code has been deposited on GitHub (https://github.com/pivlab/htp_plier). Any additional information required to reanalyze the data reported in this paper is available from the lead contact upon request.

## Acknowledgements

This work is supported by the National Human Genome Research Institute (R00 HG011898 to M.P.), and The Eunice Kennedy Shriver National Institute of Child Health and Human Development (R01 HD109765 to M.P.).

